# Differences in cell-associated and cell-free microbial DNA in blood

**DOI:** 10.1101/2025.02.13.638214

**Authors:** Kate R. Bowie, Jared Fischer, Lisa Karstens

## Abstract

In the absence of infection, blood has previously been understood to be free of microbes. However, with advances in sequencing technology this notion has been challenged, prompting new investigations into microbial DNA within the blood of both healthy and diseased individuals. To comprehensively survey microbial DNA in blood, we separated blood into fractions (plasma, red blood cells, and buffy coat) and assessed if the microbial-DNA is cell-free by the addition of DNase to a subset of each fraction. We measured 16S rRNA gene copy number with digital droplet PCR and identified the taxonomic origin of the microbial DNA with synthetic full-length 16S rRNA gene sequencing. As a use case, we examine microbial DNA from the blood of 5 men without prostate cancer (PC), 5 men with low-grade PC, and 5 men with high-grade PC. Our study demonstrates that the majority of microbial DNA is cell-free, indicating that it is not representative of proliferating microbes. Our analyses also revealed buffy coat had the lowest number of 16S rRNA gene copies yet highest number of genera of the fractions (median 23.3 copies/µL and 10 genera) and thus may be a useful fraction to study moving forward. Additionally, microbial DNA in blood may have utility as a biomarker, as we detected disease-associated compositional differences in the plasma and buffy coat fractions. This study lays the groundwork for rigorously studying microbial DNA in blood, however larger studies are needed to confirm our disease-association findings.

**Importance:** The concept of a “blood microbiome” has sparked debate in recent years, with questions about whether microbes truly exist in circulation. This study provides a crucial evaluation of the fractions of blood and their capacity to harbor microbial DNA, offering important context for prior and future research. By using DNase to differentiate between cell-associated and cell-free microbial DNA, we show that while microbial DNA is present in blood, it is sparse and heterogenous. These findings highlight the need for rigorous study design that carefully considers both positive and negative controls, as well as the specific blood fractions being examined.

## Introduction

Blood has long been considered sterile, or free of living and proliferating microbes. Recently, the potential existence of circulating microbes and their DNA in blood and its link to health and disease has garnered considerable interest. This new interest is in part due to advances in sequencing technology allowing researchers to interrogate low-microbial biomass samples, like blood, more reliably.^1–3^ Despite progress in sequencing and bioinformatic methods, microbes in blood remain controversial due to the lack of consensus on how best to handle low-microbial biomass samples and the high levels of contamination in such samples.^4^ Although controversial, some believe the microbes found in circulation are part of a circulating microbiome,^5^ while other researchers consider the microbes in circulation to be transitory.^6^ Regardless, researchers are uncovering associations between circulating microbial DNA in blood and aspects of human health with potential clinical implications, such as being predictive of response to chemotherapy treatments.^7,8^

In relatively healthy populations, studies have shown up to 100% of samples having microbial DNA in blood while others determine as few as 16% of human blood samples have microbes.^6,9,10^ This variation is likely due to differences in experimental approaches in sequencing, sample processing, and decontamination procedures. Although studies have pointed to the presence of *Staphylococcus* and *Cutibacterium*, there has not been agreement in specific bacteria found in the blood of healthy individuals indicating significant compositional heterogeneity.^6^ When it comes to evaluating microbes and microbial DNA in blood in individuals with disease, researchers have identified associations between specific genera found in circulation and hypertension as well as type 2 Diabetes.^11,12^ There have also been associations between both abundance and diversity of the microbes in blood from individuals with liver cirrhosis, respiratory diseases, and chronic kidney disease.^13–15^

In the context of cancer, approximately 16% of newly diagnosed cancers are caused by infectious agents, such as microbes.^16^ In fact, bacteria have been found in tumors themselves,^17^ and the translocation of bacteria into the blood stream has been studied with respect to colorectal cancer.^18^ In prostate cancer (PC), bacteria have been specifically implicated in inflammation of the prostate, which is important for the development of PC.^19^ To build on this work and investigate the potential of microbial DNA in blood, we designed a study to rigorously evaluate microbial DNA from blood in men with and without PC.

Overall, little is known about microbial DNA in blood, so we first set out to uncover the localization of microbial DNA in blood. The majority of prior studies have relied on plasma as a proxy for blood, even though other components of blood also have microbial DNA.^9,20^ Few studies have investigated other blood fractions (buffy coat and red blood cells),^21^ and none have investigated these fractions in disease. To build and improve on the knowledge of microbial DNA in blood, we measured the microbial load in each fraction treated with and without DNase, in non-cancer and prostate cancer patients. Additionally, many studies have used 16S rRNA amplicon gene sequencing, which targets a small region of the 16S rRNA gene. We evaluated the microbial composition using synthetic long-read sequencing of the full-length 16S rRNA gene, which increases signal to noise in low microbial biomass studies and has improved taxonomic resolution at the genus and species levels.^22^

## Results

### Study Participants and Sequencing Overview

Blood samples were taken from 15 men before undergoing prostate biopsy at Oregon Health & Science University in Portland, OR USA (under IRB# 18048 at Oregon Health & Science University, and IRB# 4214 at Portland VA Medical Center). Age and PSA were recorded for all men (**Figure S1a**). The average age of participants was 65.8 ± 5.9 years-old, with an average BMI of 28.8 ± 4.2 kg/m^2^, and average PSA of 9.2 ± 13.9 ng/mL. There were no significant differences in age, BMI, nor PSA found between disease status as determined by a Kruskal-Wallis test. The blood samples were processed to produce plasma, red blood cell pellet (RBC), buffy coat layer (BF) fractions, and one aliquot of whole blood was retained (WB). Copies of 16S rRNA genes were measured with droplet-digital PCR (ddPCR) and the remaining sample was sent to Loop Genomics for synthetic full-length 16S rRNA gene sequencing (**Figure S1b**).

### DNase-Treated Blood Samples Indicate the Presence of Cell-Free Microbial DNA in Blood

First, we measured the copies of 16S rRNA genes in each sample. The samples were normalized to volume of whole blood used to produce each fraction. Surprisingly, BF had the lowest copies of 16S rRNA genes with a median of 29.3 copies/µL (interquartile range [IQR] 17.5, 50.2 copies/µL) with plasma having slightly more (median 29.5, IQR: 27.0, 31.3 copies/µL, **Figure 1a**). RBCs followed (median 49.3, IQR: 38.7, 67.4 copies/µL), and as expected, WB had the highest copies of 16S rRNA genes (median 172.9, IQR: 133.8, 200.5 copies/µL). The amount of 16S genes in each fraction was significantly different (p-value = 4.75e-07 Friedman test). These results suggest that WB might be the best sample type for looking at microbial DNA levels.

**Figure 1.**
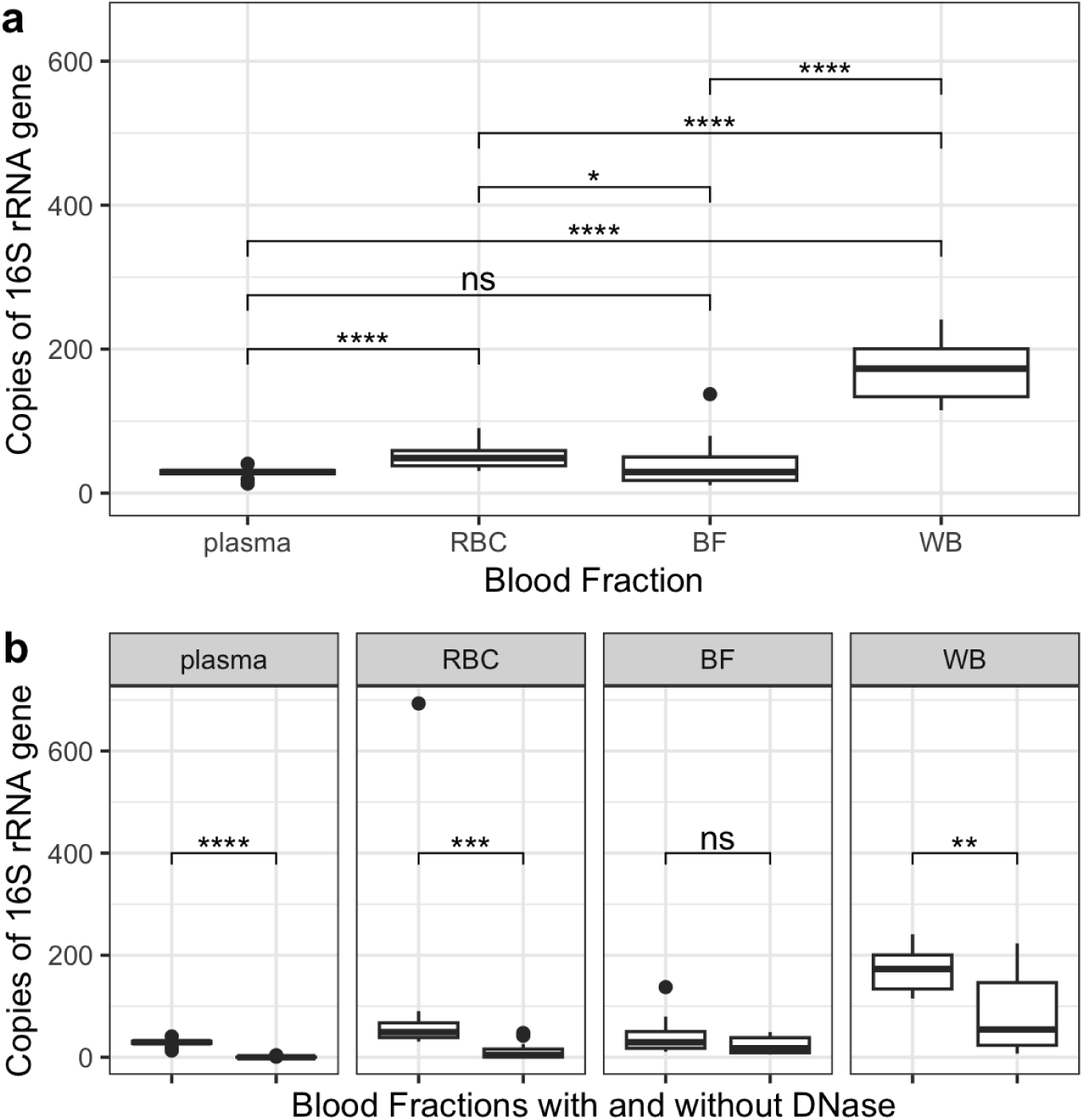
Evaluation of microbial load in blood. **a**) Boxplot displaying the number of copies of 16S rRNA genes per µL across blood fraction types. The blood fractions had significantly different number of copies determined by Friedman test (p-value = 4.75e-07). **b**) Boxplot comparing each fraction to its associated DNase-treated counterpart. All fractions had significant differences in copies/µL of 16S when compared to DNase samples.

To identify if the microbial DNA in each fraction was intracellular or extracellular, we compared each fraction with and without DNase added (**Figure 1b**). The DNase should remove extracellular DNA and leave only the intracellular microbial DNA to be measured by ddPCR. As plasma contains only cell-free DNA, it should have no copies of 16S rRNA genes when DNase has been added. As expected, the DNase-treated plasma samples had a median of 0.0 copies/µL of 16S (**Figure 1b**). The RBC DNase samples had the second lowest number of 16S rRNA gene copies (median 4.4 copies/µL, IQR: 0.5, 15.9 copies/µL). After plasma and RBC, the BF DNase samples had a median of 17.5 copies/µL (IQR: 9.0, 38.3 copies/ µL), and lastly WB had the most copies of 16S after DNase treatment with a median of 54.4 copies/µL of 16S rRNA (IQR: 23.4, 146.3 copies/µL). All the DNase samples had significantly less copies of 16S than their non-DNase counterparts (p < 0.005, **Figure 1b-c**), which indicates the presence of cell-free microbial DNA in every blood fraction.

### Negative Controls Look Distinct from Blood Fractions

After establishing the presence and quantity of microbial DNA in the blood fractions, we evaluated the composition of the samples using 16S rRNA gene sequencing. All blood fraction samples generated sequences, with a median sequencing depth of 341 reads (range 14-1,877 reads). We first examined the positive and negative controls to assess confidence in these low-microbial biomass sample sequencing results. The positive controls were mock microbial dilution series that were processed and sequenced along with the blood fraction samples (**Figure S2**). Sequencing revealed all eight expected genera in the positive controls regardless of dilution. The average percentage of unexpected taxa detected in the positive controls was 0.025% (range 0%-1.27%). The median sequencing depth of the mock community samples is 4,356.5 reads (IQR 2670 reads, 5751 reads).

Negative controls were microbial-free water blanks that were processed in triplicate alongside the blood fraction samples. The three negative controls per fraction were combined, generating an average of 864 reads (minimum 41 reads, maximum 2950 reads). The five most abundant genera in our negative controls were *Bacillus*, *Legionella*, *Prevotellaceae*, *Prauserella*, and *Escherichia-Shigella*, of which *Bacillus*, *Legionella* and *Escherichia* have previously been described as contaminating sequences in microbiome studies.^23^ The top five most abundant species in the negative controls were *Lactobacillus fermentum*, *Bacillus mycoides*, *Rubrobacter bracarensis*, *Pseudomonas lutea*, and *Methylobacterium adhaesivum*. To determine if the genera associated with the negative controls should be removed, we examined how prevalent these genera were in the associated fraction (**Figure 2a**). Outside of *Prauserella* and *Rubrobacter*, we did not identify substantial overlap between the taxa found in the samples and those of the negative controls. *Prauserella* was present in 56.3% of samples and in 75.0% of negatives (**Figure 2b**). As *Prauserella* has only been reported to be found in lakes and sediment,^24^ we removed all sequences mapping to *Prauserella*. Similarly, sequences mapping to *Rubrobacter* were present in both negative controls and samples and were removed from the data. *Nitrosomonas*, *Rhodospirillum*, *Rhodobacter*, and *Salinispora* were also removed as they have not been previously found to be associated with the human body.^25–27^

**Figure 2.**
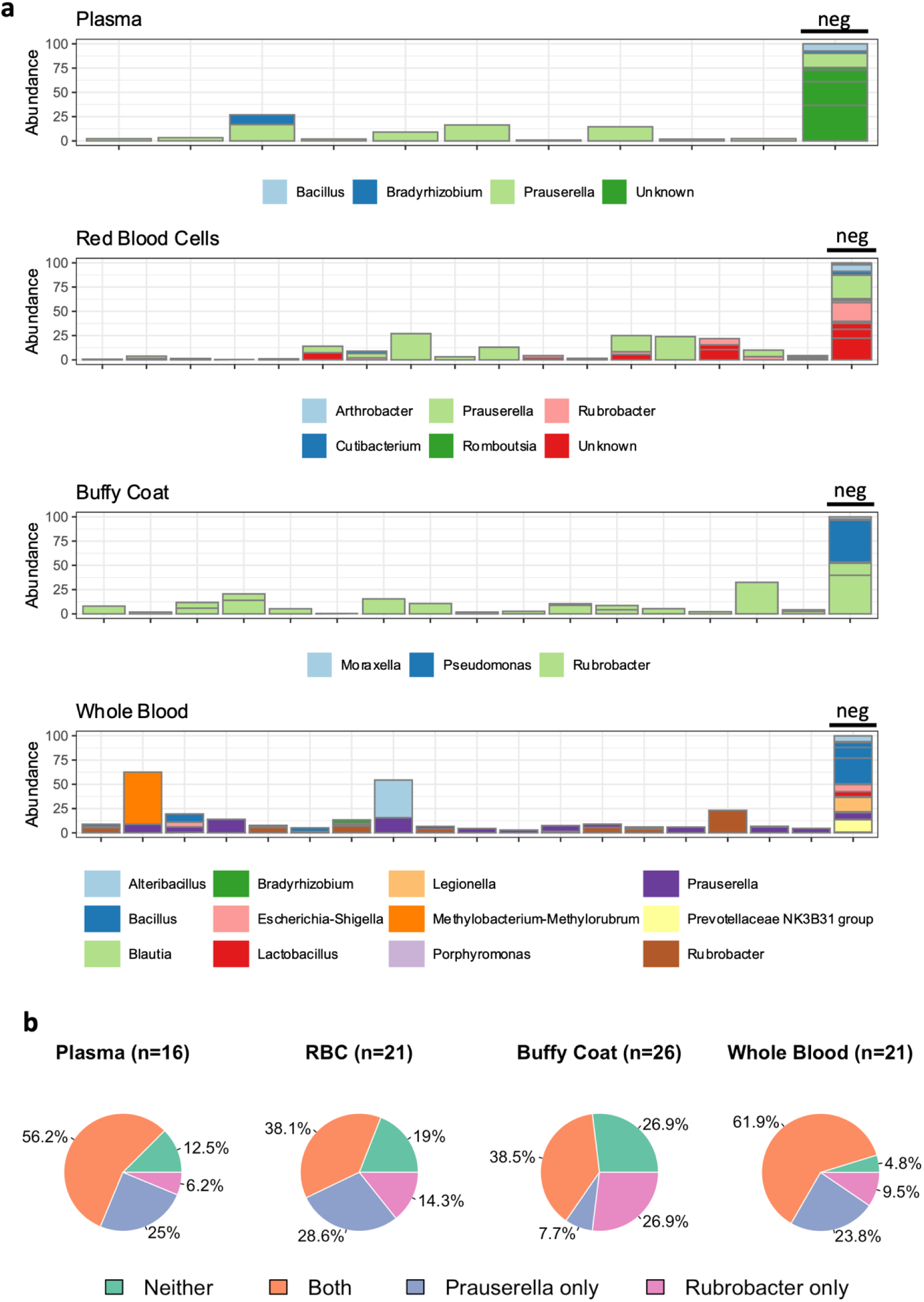
Bacteria from negative controls across samples. **a**) Stacked bar plots of the abundance of genera for each sample. All genera selected for these plots were based on the negative controls for the associated blood fraction. **b**) Pie charts demonstrating that the majority of samples had *Prauserella*, *Rubrobacter*, or both – which are considered contaminant genera in this dataset.

### Genera are Shared Across Blood Fractions

After removal of contaminants and low-quality sequences, we found there were significantly different numbers of reads for each fraction (p = 0.003, **Figure S3a**). WB had the least number of reads (median 85 reads, IQR: 59, 140 reads), followed by RBC, plasma, and finally BF with the highest number of reads (**Figure S3b**, RBC: median 166 IQR: 56, 253 reads; plasma: median 290 reads, IQR: 185, 393 reads, and BF: median 296 IQR: 196, 389). All samples were rarefied to 50 reads, which removed 20.0% of plasma and BF samples, 46.7% of RBC samples, and 66.7% of WB samples. No association was found with health status and successfully sequenced samples regardless of fraction. The sequencing data revealed 10 phylum, 92 genera and 134 species present in the blood samples, however only 82 genera and 69 species were assigned. Compositionally, Firmicutes was the most abundant phylum followed by Proteobacteria, Actinobacteria, and Bacteroidota (**Figure 3a**). The most abundant genera for all fractions were *Escherichia-Shigella, Bacillus, Bifidobacterium, Bacteroides,* and *Alteribacillus*, which was previously part of the *Bacillus* genus (**Figure S4**). All have previous been found in the human body.^28–30^ In terms of individual fractions, there was significant overlap of the top five genera, shown in **Figure 3b**. One of the most abundant genera was not assigned using our pipeline, (NA, **Figure 3b**) however we used NCBI BLAST and identified the unknown ASV as *Virgibacillus*. *Virgibacillus* has recently been found in stool samples of both children and adults.^31–33^ The most abundant assigned species were *Bifidobacterium adolescentis, Escherichia coli, Bacteroides thetaiotaomicron, Methylobacterium-Methylorubrum adhaesivum*, and *Bacillus virus*. Interestingly, only 51.5% of the taxa were assigned at the species level, and the rest could not be assigned, hence we continue the remaining analysis at the genus level.

**Figure 3.**
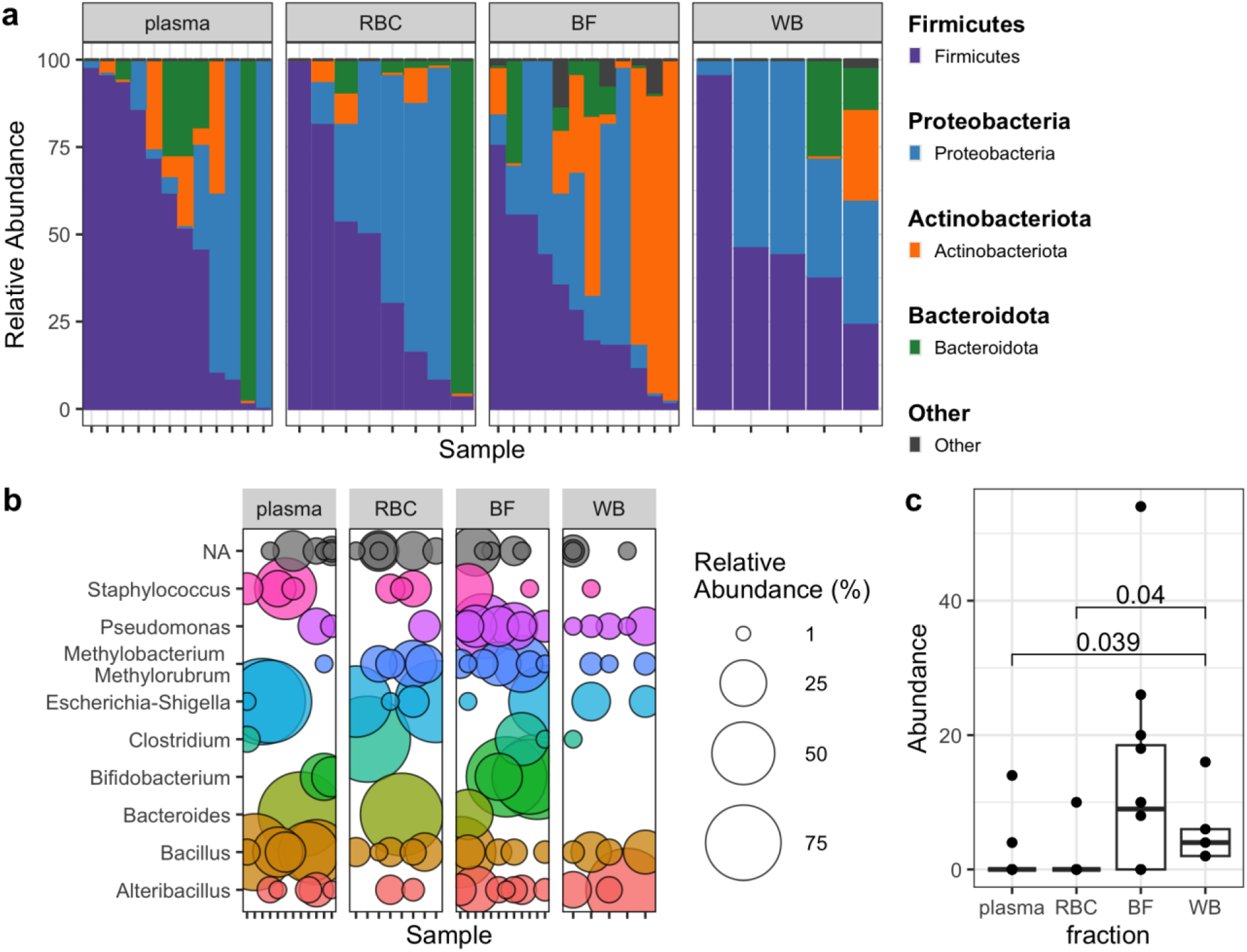
The blood fractions share similar bacteria. **a**) Firmicutes (purple) is the most abundant phylum in each of the blood fractions, followed by Proteobacteria (blue). **b**) Bubble plot of the top 10 genera separated by blood fraction. The fractions share the majority of the same genera and the bubbles display the relative abundance of each genus. **c**) Boxplots of the relative abundance of pseudomonas. WB has significantly higher relative abundance of pseudomonas compared to plasma and RBC, which have almost zero (p<0.05).

Many of the studies examining microbial DNA in blood use plasma or serum, therefore we investigated how the composition and alpha diversity of plasma differs compared to the other fractions. Plasma had the least observed genera (median 5.5 genera, range 2-13 genera) while BF had the most genera of the fractions (median 10.0 genera, range 2-14 genera), and expectedly, WB had the most observed genera of all samples with a median of 13.0 genera (range 4-17 genera, **Figure S5**). None of the alpha diversity metrics we evaluated (observed genera, Shannon index, and inverse Simpson index, Pielou) were significantly different between blood fractions. We did however evaluate the abundances of the top genera in blood and found that *Pseudomonas* was significantly different between plasma and WB, as well as between RBC and WB (p<0.05, **Figure 3c**). Next, to identify whether fractions are more similar to each other or to the patient they came from, we performed beta diversity analysis. We did not find any significant associations between any beta diversity metric (weighted UNIFRAC, unweighted UNIFRAC, and Bray-Curtis) and patient nor fraction (**Table S1**).

### Composition of DNase-Treated Samples Reveal Majority of Microbial DNA in Blood is Cell-Free

Little is known about where microbial DNA in blood is located. We sought to understand if there are compositional differences between intracellular versus extracellular microbial DNA, therefore we added DNase to samples to remove extracellular DNA. Only 54.9% of the DNase samples were successfully sequenced (**Figure S6**), whereas 100% of samples without DNase successfully generated reads. The DNase samples had significantly fewer reads than those without DNase (DNase: median 122 reads, Untreated samples: 189 reads, p = 0.045 **Figure 4a**), which is expected as the samples had less overall microbial DNA (**Figure 2a**). By number of reads, the DNase samples seemed to be largely predominated by a few genera (**Figure 4c**), with the top five being *Clostridium, Bacteroides, Escherichia-Shigella, Bacillus,* and *Bifidobacterium.* Although not statistically significant, Clostridium tended to have higher relative abundance in DNase treated samples than samples without DNase, suggesting a potential mechanism for intracellular translocation (p = 0.08, **Figure S6a**). Alternatively, Pseudomonas had a higher relative abundance in the untreated samples compared to those with DNase (p = 0.07, **Figure S6b**). We next measured alpha diversity and identified a significant difference between the diversity of samples and DNase samples (**Figure 4b**). DNase samples had a median of 3 observed genera (range 1-16) while the samples without DNase had a median of 10 observed genera (range 1-19) – a trend that held true for the Shannon and inverse Simpson indices (**Figure 4b**). These results suggest that the majority of circulating microbial DNA is extracellular.

**Figure 4.**
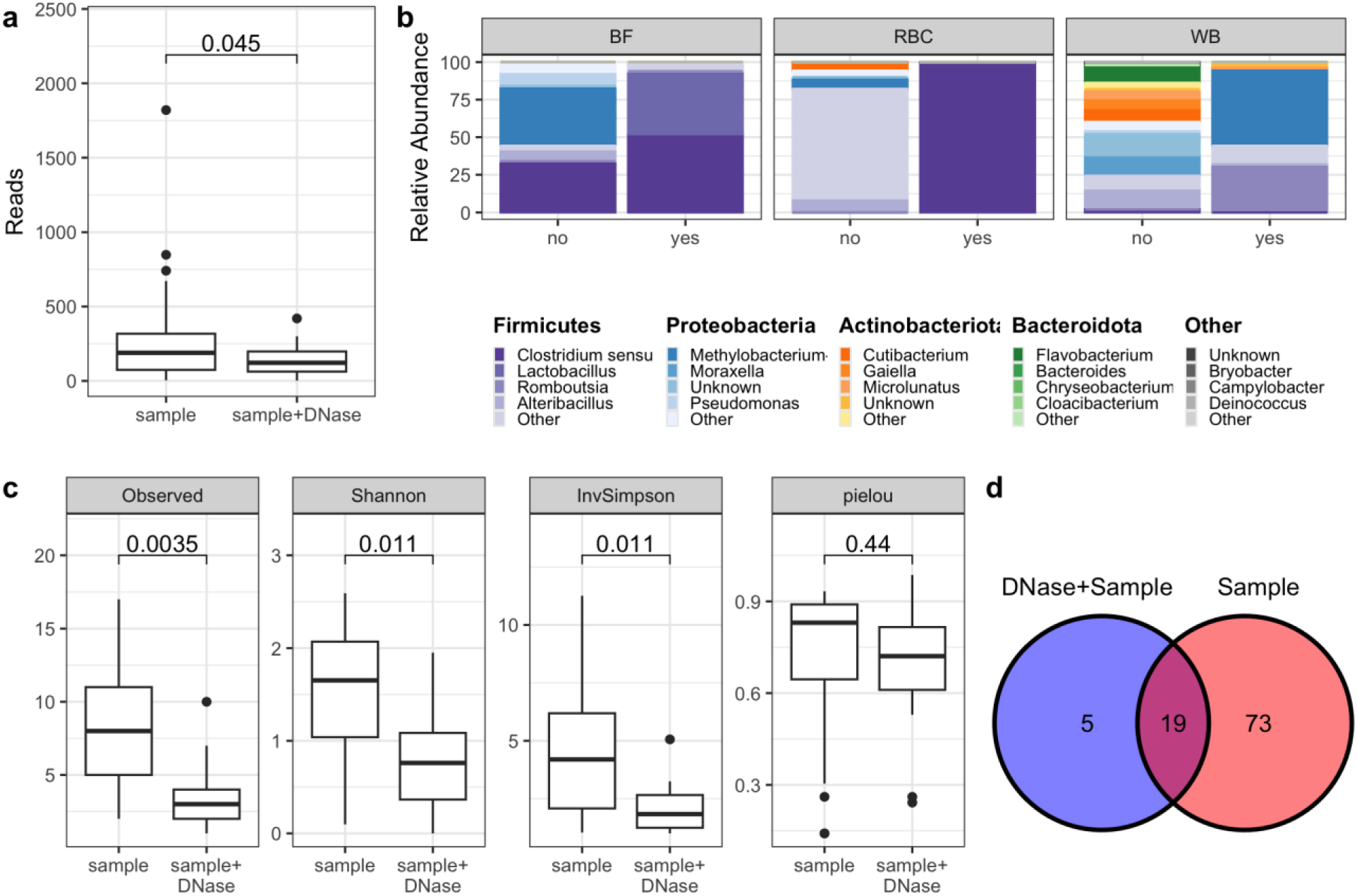
DNase blood samples have less diversity than non-DNase samples. **a**) Boxplot displaying the significant difference in sequencing reads between DNase and non-DNase samples (p= 0.045). **b**) Stacked bar plot of microbial DNA found in paired DNase and non-DNase blood fractions for patients 10, 14, and 4. **c**) Summary of alpha diversity metrics between DNase and non-DNase samples, demonstrating the significant decrease in alpha diversity in DNase samples. **d**) Venn diagram indicating the number of distinct and shared taxa between DNase and non-DNase samples.

We also evaluated beta diversity metrics of samples with and without DNase and found significant associations with both unweighted UNIFRAC and the Bray-Curtis beta diversity metrics (p = 0.02 and p = 0.04 respectively, **Figure S7b-c**). To better understand which taxa were potentially contributing to these beta diversity dynamics, we examined the genera present only in the DNase samples and vice versa. We identified 5 genera present only in the DNase samples, however 2 genera were not assigned at the Genus level. Using NCBI BLAST, we identified one sequence as *Finegoldia magna*, an opportunistic human pathogen often found in the human gut, urinary tract, and skin (**Figure 4e**).^34–36^ Conversely, we identified 73 genera found only in the non-DNase samples, meaning these genera are only located extracellularly in this patient population. 19 genera were shared between DNase and non-DNase samples, which could hint at taxa that can translocate.

### Both the Plasma and Buffy Coat Fractions have Evidence of Disease-specific Associations with Beta Diversity

Next, we examined if there are any disease-associated patterns in microbial DNA in blood. Our patient population consists of 15 men, made up of three groups of five men each. The groups are composed of individuals who are screen-negative for PC, low-grade PC, and high-grade PC. Given the low sample numbers for the RBC and WB fractions (**Table S2**), we focused our PC analysis on plasma and BF. We first evaluated alpha diversity between the disease groups and did not find any associations (**Figure S8**). We did however identify significant associations between both weighted UNIFRAC and disease when adjusting for fraction (weighted UNIFRAC: p = 0.008, **Figure 5a, top**), as well as Bray-Curtis and disease (p = 0.03). This indicates that there are compositional differences in the circulating microbial DNA between men that are screen-negative, diagnosed with low-grade PC, and diagnosed with high-grade PC taking into account blood fraction. Next, we created a biplot at the phylum level to begin to uncover which taxa could be responsible for the compositional difference between disease states (**Figure 5a, bottom**). This biplot, along with an overview of the data (**Figure S9**), demonstrated that many of the high-grade PC samples had a high abundance of Proteobacteria, which could potentially be an indicator of more severe disease. We next visualized the mean relative abundance of the top four phyla (Bacteroidota, Actinobacteriota, Proteobacteria, and Firmicutes) across the disease groups in plasma or buffy coat with a chord diagram (**Figure 5b**). Although we are likely underpowered, it is clear that the distribution of phyla differs depending on fraction, yet they both show distinct patterns which could potentially distinguish high-grade PC from low-grade PC and screen-negative patients. For example, while both plasma and buffy coat high-grade PC samples demonstrate high abundance of Proteobacteria (**Figure 5a**, **Figure 5c** p=0.17, **Figure 5d** p=0.09), only the low-grade plasma samples are predominated by Firmicutes (**Figure 5b**, **Figure 5d**). Although it did not withstand multiple test correction, we also found the buffy coat fraction demonstrated a higher abundance of Actinobacteriota in the screen-negative samples as compared to the other disease groups (unadjusted p-value=0.08, adjusted p-value=0.17 **Figure 5d**). At the genus level, we did discover trends with a genus of the Proteobacteria phyla, *Methylobacterium-Methylorubrum* (**Figure S10**, p=0.07), which was not present in screen-negative patients. This could be a result of an impaired immune system, as various species of *Methylobacterium* have been found to cause hospital-acquired infections in previous literature.^37^ Additionally, we examined associations between alpha diversity and PSA, BMI, and age, but did not find any significant associations.

**Figure 5.**
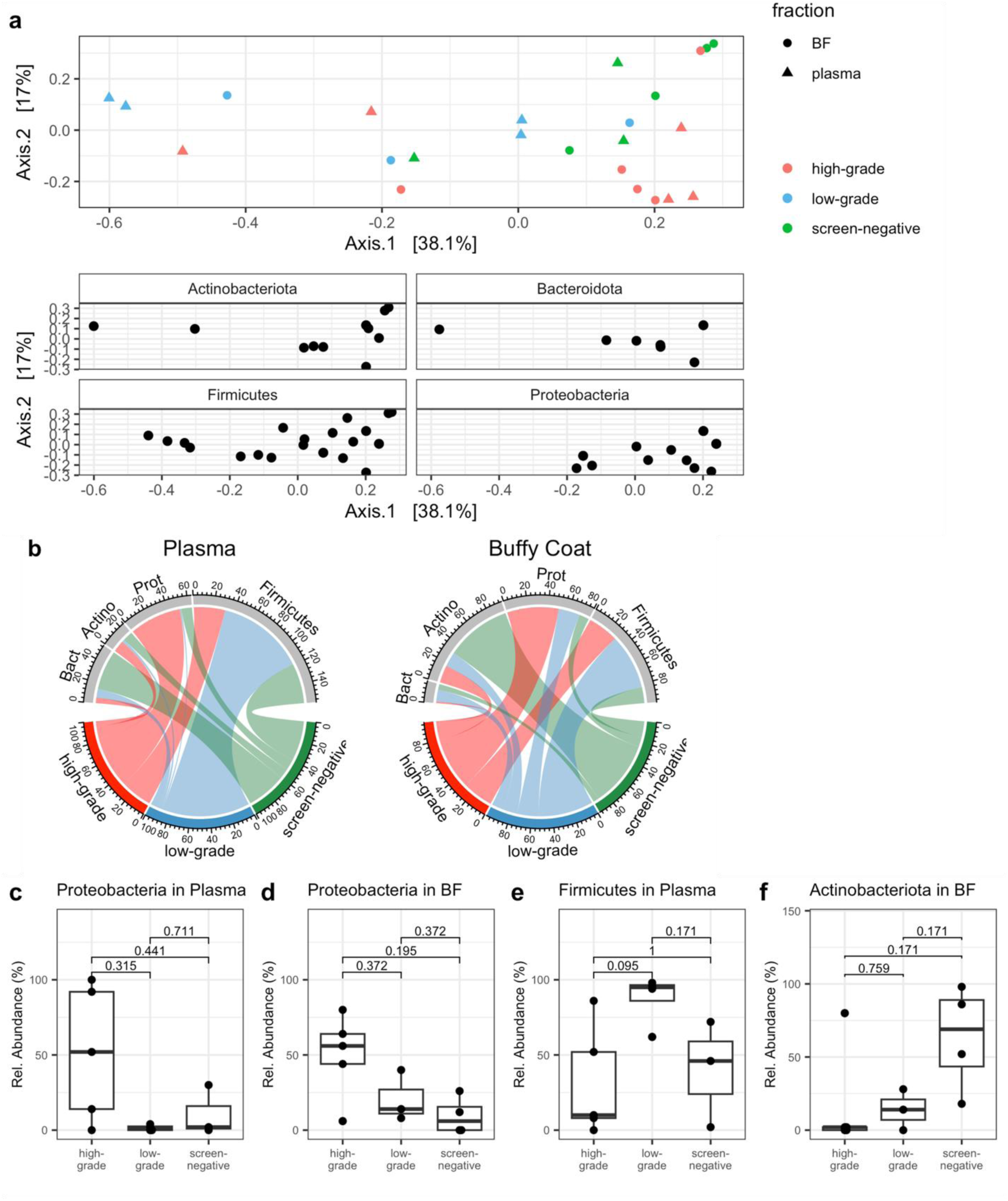
Phylum-specific beta diversity associations with disease depend on fraction. **a**) (Top) Weighted UNIFRAC principal coordinate analysis (PCoA) plot of the plasma (triangle) and buffy coat (circle) fractions colored by disease status (p=0.008). (Bottom) Biplot demonstrating the phyla associated with each sample’s composition. **b**) Chord diagrams at the phylum level for plasma and buffy coat samples demonstrating the average relative abundance of each phylum (Bact=Bacteroidota, Actino=Actinobacteriota, Prot=Proteobacteria) in each disease category. **c**) Boxplots demonstrating the relative abundance of Proteobacteria in plasma and **d**) buffy coat, **e**) Firmicutes (plasma only), **f**) Actinobacteriota (buffy coat only) in each disease group.

## Discussion

The results from our study indicate that there is measurable microbial DNA in circulation. Using full-length 16S rRNA gene sequencing, we identified 78 genera and 69 species with high confidence. In a similar study, Tan et al found 56 genera and 117 species, however, this study had a much larger number of participants (10,000) and used a different sequencing modality and decontamination parameters which could explain the disparity in number of taxa.^6^ This discrepancy points to the importance of sequencing and data processing and how they can impact the resulting data.

Through our early exploratory data analysis, we noticed a stark difference between the number of sequencing reads and number of copies for the WB samples. Specifically, WB had the highest microbial load measured by copies of 16S (**Figure 1a**), yet the lowest number of reads. We speculate that the WB samples have a high number of short sequencing reads (<1350 bp) as compared to the other fractions, which were removed during data processing. This potentially points to WB having fragmented microbial DNA, which can be measured with ddPCR yet would produce a low number of reads from full-length 16S gene sequencing.

BF includes white blood cells which are understood to engage with microbes as part of a normal immune response,^38^ therefore we expected BF to have more associated microbial DNA resulting in the highest 16S rRNA gene copy number and number of genera. Instead, we found that BF contains the least amount of microbial DNA (**Figure 1a**), although plasma had a lower mean (28.4 copies/µL versus BF at 41.5 copies/µL) and had the fewest number of genera (**Figure S5**). Our results also demonstrated that the RBC fraction has more copies than BF (**Figure 1a**). These results are contrary to a previous study, although we utilized ddPCR which is known to produce more precise and reproducible results as compared to qPCR.^9,39^

Unexpectedly, we discovered the RBC fraction to have a similar number of observed genera to BF (median 9.5 genera compared to 10 genera, **Figure S5**). These findings suggest an interaction between RBCs and circulating or transitory microbes. In fact, in conditions of bacteriemia, RBCs have been documented to interact with bacteria by either entrapping, killing, then releasing the dead bacterium back into circulation,^40^ or the RBCs use their electric charge to attract and kill bacteria.^41^ Our findings could point to similar mechanisms of RBCs as seen in bacteremia. Not only this, but RBCs have a life cycle of approximately 120 days, thus studying the RBC fraction for microbial DNA could serve as a recent history of the types of microbes that have translocated in the body.^42^

A major finding of our study is that the majority of microbial DNA in blood is cell-free. No other studies have used DNase to explore the question of intra-versus extracellular microbial DNA. We found the DNase-treated samples to have significantly less copies of 16S per µL (**Figure 1b-c**) and lower complexity (**Figure 4b-c**). We did however discover the DNase-treated samples shared 79.2% of their genera with the non-DNase treated samples (**Figure 4d**) and tended to have a higher relative abundance of *Clostridium* (**Figure S6**). We used NCBI Blast and found the *Clostridium* sequence to map to *Clostridium sporogenes* – which is a spore-forming bacteria and while uncommon, has previously been documented to cause bacteremia.^43^ *C. sporogenes* is a member of the normal gut microbiome, however in the spore state, *C. sporogenes* are dormant and can survive in hostile environments even in the absence of nutrients,^44^ perhaps allowing it to go undetected in blood. Many of our DNase-treated samples did not successfully sequence, which inhibited the analysis we could perform. Future studies in this area should include examining how disease potentially affects the composition in DNase-treated samples, which could indicate changes to the cells that make up the RBC and BF fractions.

Supporting prior research of circulating microbial DNA associations with disease, we did determine compositional differences in the plasma and buffy coat fractions between PC-negative samples as well as those with low- and high-grade disease. With a larger sample size, a more robust analysis of the alpha diversity differences in specific fractions with respect to disease could give us insight into how disease impacts these transitory microbes. Specifically, more samples are needed to study the potential trend of Proteobacteria being more abundant in high-grade PC samples (**Figure 5c**). What is unclear is if the patterns we discovered are a general cancer-related finding, or specific to PC.

A limitation of this study was the small sample size, which was a tradeoff we made to increase rigor. As we processed every fraction in triplicate and doubled that to treat samples with DNase, we processed a total of 360 samples not including the controls. We believe the replicates are important for capturing the diversity of these low-microbial biomass samples, however a small study of 15 patients quickly became cumbersome. Full-length 16S rRNA gene sequencing gave us high confidence in the resulting taxonomy and allowed us to rarefy to a lower read-depth than traditional 16S amplicon sequencing.^3^

In summary, our study indicates the presence of circulating microbial DNA in both PC negative and positive men. Our results suggest microbial DNA in blood is sparse, heterogenous, yet present in all fractions of blood. We revealed that not only is the majority of microbial DNA in blood cell-free, but also that the majority of the microbial diversity stems from the BF fraction (as opposed to plasma or RBC fraction). The sequencing results demonstrated that plasma samples had the most samples successfully sequence while WB had the least which can inform future study design. Lastly, although we saw trends in the plasma and BF fractions of our PC cohort, more studies with larger sample sizes are needed to further investigate associations with specific fractions of blood and disease.

## Methods

### Patient Cohort

Our research complies with all relevant ethical regulations of Knight Cancer Institute at Oregon Health & Science University. Patients were undergoing prostate biopsy and were consented through VAPORHCS/OHSU: Cancer Early Detection Advanced Research (CEDAR) Specimen and Data Repository. The associated IRB# is 18048 at Oregon Health & Science University, and IRB# 4214 at Portland VA Medical Center. Participants were not compensated.

### DNA Isolation and Sequencing

Blood was drawn from men with high-grade PC, low-grade PC, and without cancer (n=5 per group). Samples were processed into the plasma, buffy coat (BF), red blood cell pellet (RBC) and whole blood (WB) fractions in six 200 µL aliquot per fraction prior to freezing. For each sample, 5 mL of blood was removed from the tube and 600 µL whole blood was removed, followed by a room temperature 1200 rpm centrifugation step for the plasma fraction. The remaining 8-10 mL underwent density gradient centrifugation using Ficoll-Paque PLUS (Cytiva) for the buffy coat and red blood cell pellet fractions. To examine if microbial DNA was extracellularly or intracellularly located, DNase (New England Biosciences) was added to three of the six replicates for each fraction, and the product protocol used. Once all blood fractions were aliquoted into three replicates and DNase added to the remaining three, samples were frozen. Microbial-free water (Qiagen) was used as negative controls and a mock microbial community (Zymo) was serially diluted,^4^ and processed alongside blood samples. Microbial DNA was extracted using the QiAMP DNA mini kit (Qiagen). Extracted DNA from the blood fractions was submitted to Element Biosciences for their 16S rRNA synthetic long-read sequencing using LoopSeq technology.

### Droplet-Digital PCR

Droplet-digital PCR (ddPCR) was used to measure 16S rRNA copies. Each PCR reaction was prepared in a PCR hood in a dedicated ddPCR room. For the mock microbial community dilution series, each dilution was diluted either 1:1000 or 1:100 in microbial-free water (Qiagen). The QX200 Digital PCR System with Auto Droplet Generator and Reader along with the QX200 ddPCR EvaGreen Supermix (Biorad) was used. The PCR protocol uses a 2°C/second ramp rate and starts with a 10-minute 95°C enzyme activation step followed by 40 cycles of a two-step protocol (94°C for 30 seconds and 59°C for 1 minute), and lastly cycles up to 98°C for 10 minutes. V6 primers from the Sfanos lab at Hopkins were used (Forward: CAACGCGWRGAACCTTACC; Reverse: CRRCACGAGCTGACGAC). The results were then normalized to starting whole blood volume required to produce 200 µL of each blood fraction for comparison across fractions.

### Bioinformatics and Statistical Analyses

Loop Genomics provided us with assembled full-length 16S rRNA gene sequences. These sequences were processed into amplicon sequence variants (ASVs) using DADA2.^3^ The RDP Classifier was used to map the ASVs to the SILVA 138 16S rRNA reference set for taxonomic identification. All analyses were completed on sequences between 1350 and 1600 base pairs length. The mock community dilution series and negative controls were analyzed to determine how to handle biological samples using phyloseq (version 1.42.0) and visualized using microshades (version 1.11).^45^ ASVs that could not be assigned at the Family level or higher were discarded. ASVs originating from the negative controls (**Figure 2**) were removed. All subsequent analyses were done on ASVs agglomerated at the Species or Genus level in R version 4.3.2. The replicates for each sequencing sample were combined prior to rarefaction to represent one microbiome composition per fraction per individual. Rarefaction to 50 reads was based on retaining the maximum number of samples while accurately representing the composition (**Figure S11**). The *vegan* R package version 2.6.4 and *rstatix* version 0.7.2 were used for all statistical analyses. We also evaluated the relative abundance of the top 5 phyla and top 10 genera. A Kruskal-Wallis test was used to test for a significant associations between relative abundance and clinical characteristic of interest, and then a pairwise Wilcoxon Rank Sum test with FDR correction.

## Declarations

## Acknowledgements

This research is supported by NIH/NIDDK K01DK116706 (LK) and the Cancer Early Detection Advanced Research Center at Oregon Health & Science University’s Knight Cancer Institute (KB, JF, LK).

## Author contributions

KB, LK, and JF designed the study. LK designed and guided the data analysis and wrote the manuscript. KB processed all samples, analyzed the data, and wrote the manuscript. All authors contributed to interpretation of results and editing the manuscript.

## Availability of data and material

Processed data and code used to process and analyze the data is available on the Karstens github (https://github.com/KarstensLab/microbial_dna_in_blood).

